# A stop codon shared by all *Gyps* vultures, including the three Critically Endangered species of South Asia, lies in a coding exon of the bird gene *CYP2C45*

**DOI:** 10.64898/2026.11.22.768365

**Authors:** Chandrani Das, Krishnendu Sinha

## Abstract

Veterinary diclofenac devastated three South Asian *Gyps* vultures, but the basis of their sensitivity is unclear. A stop codon reported in a Cape Vulture P450 gene, described as *CYP2C19* in a retained intron, was re-examined with public data: 45 genome assemblies, raw reads of seven *Gyps* birds and a liver transcriptome. The gene is the bird *CYP2C45*, without a one-to-one human orthologue. The stop (codon 338, CGA→TGA) lies in coding exon 7; the intron is spliced normally, the mRNA carries the stop, and the truncated protein lacks the haem-binding cysteine. The stop was found in all eight living *Gyps* species, homozygous wherever testable, and in none of 27 other raptors, including two killed by diclofenac. It arose once, 1.5–10.6 million years ago on the *Gyps* stem, and the gene has since evolved near-neutrally. This genus-wide loss marks *Gyps* vulnerability but cannot explain diclofenac deaths in other raptors.

## 1 Introduction

In the 1990s the populations of three vultures of the Indian subcontinent, the White-rumped Vulture (*Gyps bengalensis*), the Indian Vulture (*G. indicus*) and the Slender-billed Vulture (*G. tenuirostris*), fell by more than 95%. The cause was diclofenac, an anti-inflammatory drug given to cattle: vultures that fed on treated carcasses died of kidney failure with visceral gout (Green et al., 2004; Oaks et al., 2004; Shultz et al., 2004). Dosing trials then showed that diclofenac is just as toxic to the African White-backed Vulture (*G. africanus*), the Griffon Vulture (*G. fulvus*) and the Cape Vulture (*G. coprotheres*) (Naidoo et al., 2009; Swan et al., 2006), and that the Turkey Vulture (*Cathartes aura*) tolerates doses more than a hundred times larger (Rattner et al., 2008). India banned veterinary diclofenac in 2006 and two related drugs, ketoprofen and aceclofenac, in 2023 (BirdLife International, 2023; Ministry of Health and Family Welfare, Government of India, 2023), but the three species remain Critically Endangered.

Why are *Gyps* vultures so sensitive? Diclofenac lingers in their bodies: it is cleared slowly, and it interferes with the transport of uric acid out of kidney cells (Adawaren et al., 2024; Naidoo and Swan, 2009). In humans, the main oxidative route for diclofenac runs through CYP2C9, a cytochrome P450 enzyme of the CYP2C subfamily (Leemann et al., 1993). Slow clearance in vultures therefore pointed to a weak or missing CYP2C enzyme. Adawaren et al. (2024) searched the Cape Vulture genome and liver transcriptome for the gene they called *CYP2C19* and found a premature stop codon, described as lying in intron 7 and retained in the mature mRNA. Colston et al. (2025) then searched public genomes and found the same stop in three more *Gyps* species (*G. africanus, G. fulvus, G. himalayensis*) but not in ten other species (nine vultures and the Steppe Eagle).

Three questions remained open. First, the three species that actually collapsed in South Asia have no genome assembly, so whether they carry the stop was unknown (Colston et al., 2025). Second, a stop codon in an intron is unusual; whether the codon really sits in an intron that is kept in the mRNA, or in an ordinary coding exon, changes how the defect should be understood. Third, birds have their own set of *CYP2C* genes that arose separately from the human ones (Watanabe et al., 2013), so whether the vulture gene is truly the counterpart of human *CYP2C19* matters when results from human pharmacology are carried over.

Here these questions are answered with public data alone. The gene is identified by its neighbours and by gene trees; the position of the stop is fixed on an exon–intron map supported by RNA-seq; the stop is read directly from raw genome reads of the three South Asian species and of four further birds, among them Rüppell’s Vulture, whose assembly lacks the codon; and 27 other raptors are surveyed, including every other vulture species with a genome assembly and species with known outcomes after diclofenac exposure. Two further questions follow from the answers: when the stop arose, and whether the gene has since evolved as a gene without function.

## 2 Results

### 2.1 The gene is the bird-specific *CYP2C45*, not an orthologue of human *CYP2C19*

The gene was located in the long-read assembly of *G. africanus* by aligning chicken cytochrome P450 proteins to the genome (Methods). It lies on contig JBEHDR010000102.1 between *ZP4* and *TLL2*, in the same neighbourhood as in the chicken: *SORBS1* –*PDLIM1* –*HELLS* –*ZP4* –[gene]– *TLL2* –*TM9SF3* –*PIK3AP1* (Fig. 1B; Table S6). The chicken gene at this position is *CYP2C45* (Watanabe et al., 2013) *(RefSeq NP_001001752*.*2; listed by NCBI under the symbol CYP2C18* with the synonym *CYP2C45*). The vulture protein is 80.8% identical to it. Chicken also carries a second, 99.4%-identical copy next to it (*CYP2C18L*, XP_015144012.2); vultures and the other raptors examined have a single copy. A second bird *CYP2C* gene, *CYP2C23* (formerly *CYP2H*), lies about 1.27 Mb away on the same vulture contig, next to *ABCG2* and *ALOX5*, again as in the chicken.

**Figure 1:**
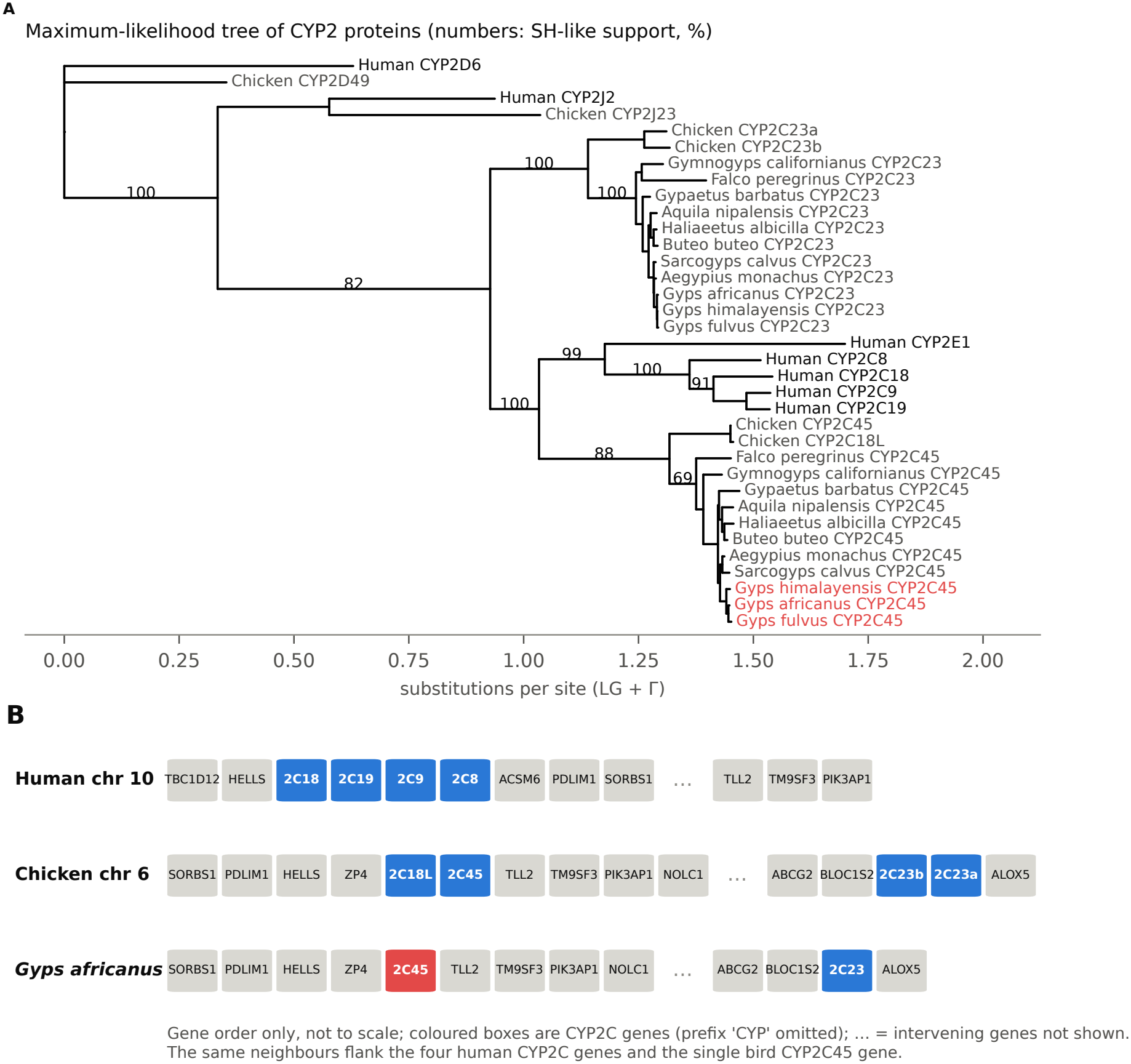
The vulture gene is *CYP2C45*, the bird counterpart of the whole human *CYP2C* cluster. **(A)** Maximum-likelihood tree of CYP2 proteins (LG + Γ; SH-like support shown for main branches). Human *CYP2C8, CYP2C9, CYP2C18* and *CYP2C19* form one group and bird *CYP2C45* another; *Gyps* proteins in red. The neighbour-joining tree is in Fig. S1. **(B)** Gene order around the *CYP2C* genes in human, chicken and *G. africanus*.

In humans, the same neighbours (*HELLS, PDLIM1, SORBS1, TLL2, TM9SF3, PIK3AP1*) surround all four human *CYP2C* genes, *CYP2C18, CYP2C19, CYP2C9* and *CYP2C8* (Fig. 1B). In a protein tree, the four human genes form one group (support 100% in both the maximum-likelihood and the neighbour-joining tree) and the bird *CYP2C45* genes form another (88% and 100%); *CYP2C45* never groups with *CYP2C19* alone (Fig. 1A; Fig. S1; Table S7). The four human genes therefore arose by duplications in the mammal line, after birds and mammals separated. The bird *CYP2C45* is the counterpart of the whole human cluster, not of *CYP2C19* in particular. The bird name, *CYP2C45*, is therefore used for the vulture gene from here on. It is the same locus that was called *CYP2C19* by Adawaren et al. (2024) and Colston et al. (2025) (Supplementary Note S1).

A wider tree confirms this (Fig. S3; Tables S15, S16). Every RefSeq protein of the CYP2C and CYP2H subfamilies was added from thirteen more vertebrates (two owls, Golden Eagle, Zebra Finch, Mallard, turkey, chicken, alligator, turtle, anole lizard, *Xenopus*, mouse and human), giving 117 proteins, and a maximum-likelihood tree was built with IQ-TREE (Methods). The eighteen mammalian CYP2C genes (four human, fourteen mouse) form one clade (SH-aLRT/ultrafast bootstrap 99.6/100), and the avian CYP2C45 genes, including those of owls, Zebra Finch, Mallard and turkey, form another (100/100), which joins two alligator genes in an archosaur *CYP2C* lineage. The two clades meet only deep in the tree. Trees forcing avian CYP2C45 to pair with any single human gene, *CYP2C8, 2C9, 2C18* or *2C19*, were all significantly worse than the unconstrained tree (approximately unbiased test, *p*-AU *≤* 0.0007; Table S15). Automated annotation names the avian orthologues inconsistently, as ‘CYP2C19’, ‘CYP2C19-like’, ‘CYP2C9’, ‘CYP2C9-like’ or, in chicken, ‘CYP2C18’ (Table S16), which probably explains how the vulture gene came to be called *CYP2C19*. The tree also shows that birds carry other *CYP2C* -like genes, for example a *CYP2C5* -like lineage in owls, Zebra Finch and Mallard; these were not examined in vultures.

### 2.2 The stop codon lies in coding exon 7, at codon 338 of 496

In *G. africanus, CYP2C45* has nine coding exons spread over 5.8 kb and a reading frame of 496 codons (Fig. 2A). Codon 338, near the start of exon 7, is TGA, a stop codon. In all non-*Gyps* raptors the same codon is CGA (arginine), except the falcon and one of the two haplotypes of a Bearded Vulture, where it is CAA (glutamine) (Fig. 2C). The change from CGA to TGA is a single C*→*T transition at a CpG site, the most mutation-prone kind of site in vertebrate genomes. The codon lines up with arginine 333 of human CYP2C9 and CYP2C19, a position conserved across the subfamily (Table S8). This is the variant reported earlier: the 66-nucleotide sequence published by Colston et al. (2025) maps to positions 8,404,695–8,404,760 of the same contig, and the codon occupies positions 8,404,731–8,404,733 (Supplementary Note S1).

**Figure 2:**
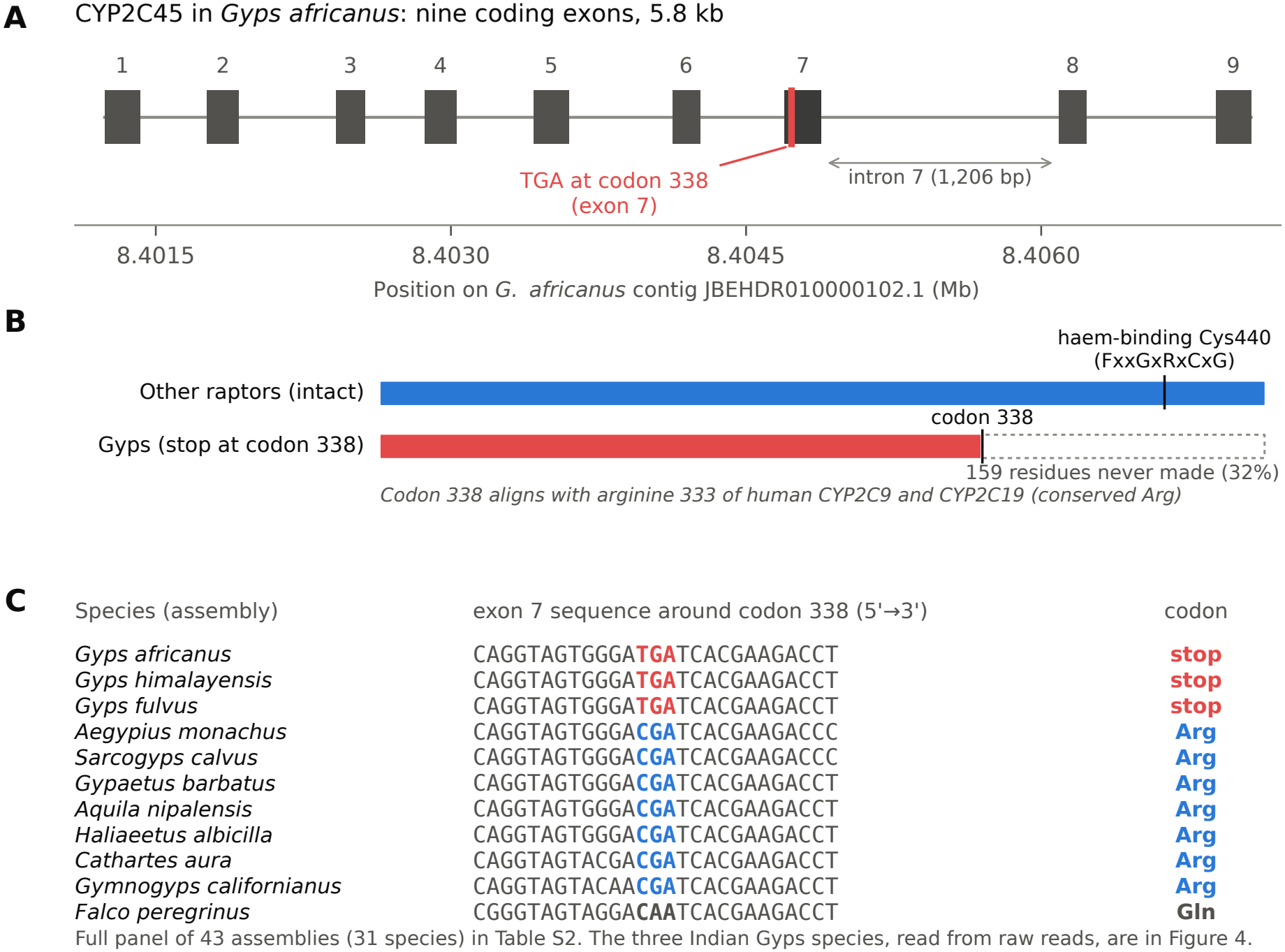
The *CYP2C45* stop codon in *Gyps*. **(A)** Gene model in *G. africanus*; boxes are coding exons. The TGA at codon 338 lies in exon 7, upstream of intron 7. **(B)** Predicted proteins. In *Gyps* the protein stops after residue 337 and lacks the last 159 residues, including the haem-binding cysteine (Cys440). **(C)** Exon 7 sequence around codon 338 in representative species (all assemblies in Table S2).

The position matters. Exon 7 is bounded by a normal AG acceptor (end of intron 6) and a normal GT donor (start of intron 7) in every assembly in which it was recovered; all introns use canonical GT–AG (or GC–AG) splice sites (Table S2). Intron 7 begins 150 nucleotides downstream of the stop codon and is 1,206 bp long in *G. africanus*. The stop therefore sits in coding sequence that every raptor uses as exon, not in an intron.

A protein that ends at residue 337 lacks its last 159 residues (32%; Fig. 2B). These include the haem-binding loop (FxxGxRxCxG, cysteine at residue 440), whose cysteine holds the iron atom at the heart of every P450 enzyme. Without it, the product cannot bind haem and cannot oxidise drugs. The defect is therefore a loss of the enzyme, not a weakening of it.

### 2.3 The liver mRNA is spliced normally and carries the stop

To see how the gene is used, a public Cape Vulture liver transcriptome (8.6 million read pairs; BioProject PRJNA912371) was mapped to the *G. africanus* contig (Methods). The two species differ little, and only alignments with at least 95% identity were kept. In all, 1,217 read pairs mapped to the two *CYP2C* genes.

Intron 7 is removed normally. Twenty-four fragments join the end of exon 7 exactly to the start of exon 8, and only a few run on into the intron, giving an intron-retention value of 0.09 (Fig. 3A,B; Table S5). This is no higher than for introns 3 to 8 as a group (0 to 0.09; median for the gene 0.06), and the intron is covered at about 3% of the depth of the exons on either side. Exon 7 is joined to exon 6 at its start (8 fragments), exactly as the gene model predicts. Thus the stop codon lies in a properly spliced exon of the mature message. All 19 mRNA fragments with a high-quality base at the first position of codon 338 read T, and all fragments at the second and third positions read G (20 of 20) and A (22 of 22) (Fig. 4B): the mature mRNA carries TGA.

**Figure 3:**
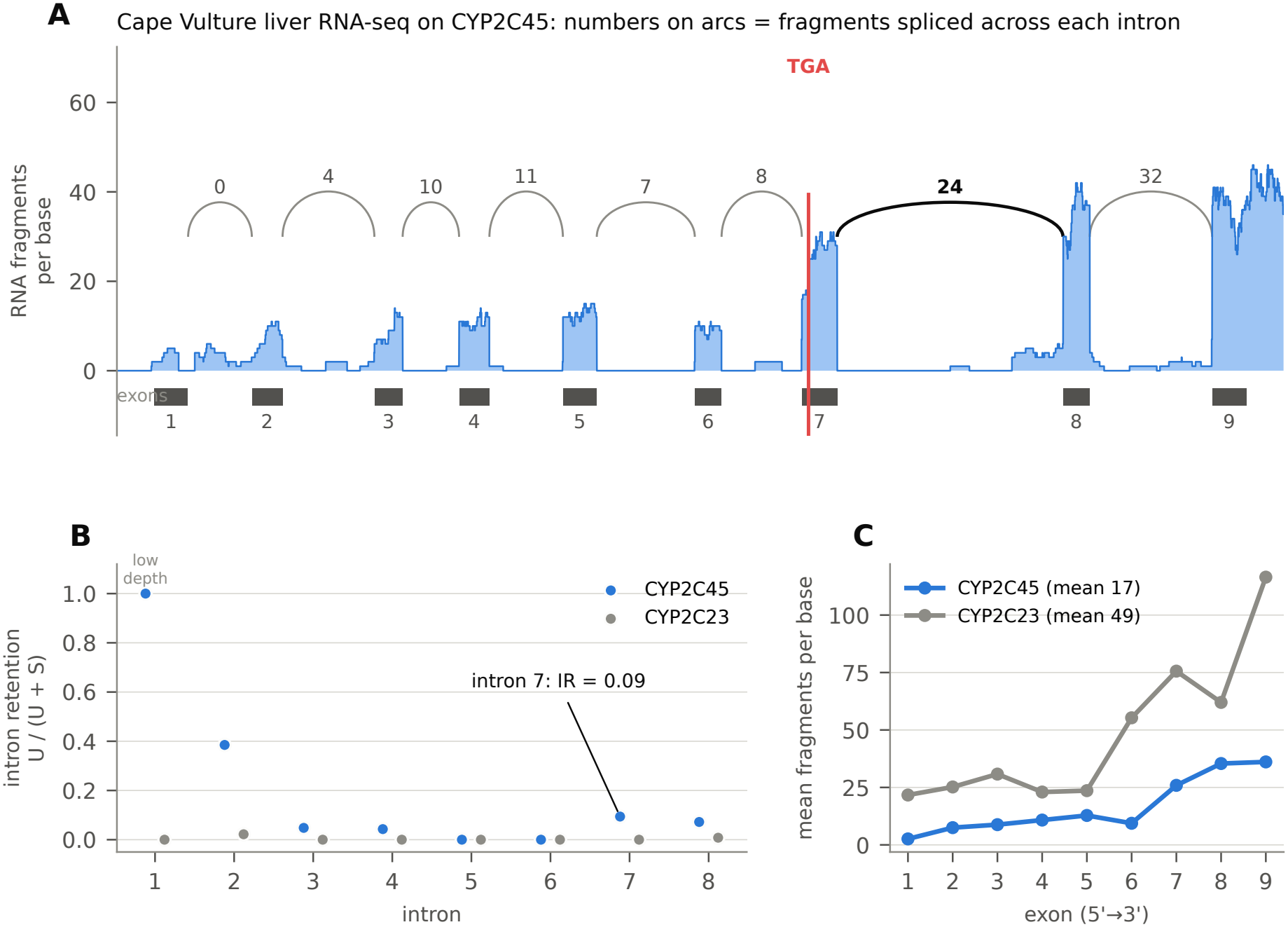
Intron 7 is spliced normally in the Cape Vulture liver. **(A)** RNA-seq coverage along *CYP2C45* ; arcs show fragments spliced across each intron (intron 7 in bold). The red line marks codon 338. **(B)** Intron retention, *U/*(*U* + *S*), for each intron of *CYP2C45* and of the neighbouring gene *CYP2C23*. Intron 1 shows no canonical splicing because an alternative donor is used (Note S4); intron 2 has few reads. **(C)** Mean coverage of each exon. Both genes show more reads towards the 3^*′*^ end, a common library effect

**Figure 4:**
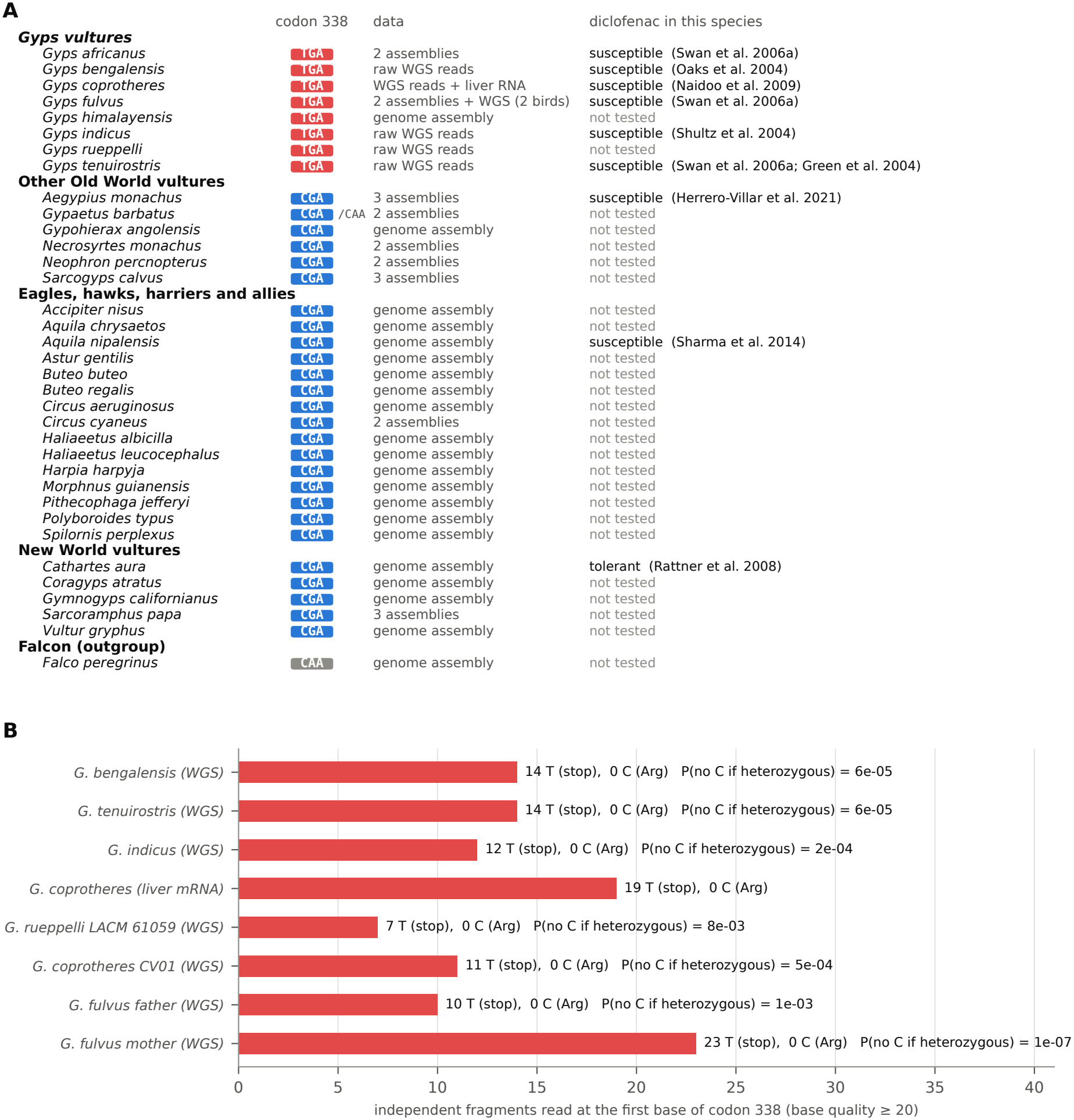
The stop is present in every *Gyps* species examined and in no other raptor. **(A)** Codon 338 in 35 species, with the data source and what is known about diclofenac in each species. Susceptible: deaths after experimental dosing or field deaths with diclofenac residues and gout. **(B)** Independent DNA fragments (raw genome reads) or mRNA fragments (Cape Vulture liver) showing T (stop) or C (arginine) at the first base of codon 338.

The gene is still expressed in the liver, at about one third of the level of its neighbour *CYP2C23*(mean depth 16.7 versus 48.9 fragments per base; Fig. 3C). The stop lies 292 nucleotides upstream of the last exon–exon junction. Messages with a stop more than 50–55 nucleotides before the last junction are normally destroyed by nonsense-mediated decay (Nagy and Maquat, 1998), so the true level of intact message is probably lower than that of the gene’s transcription. Two minor patterns were also seen: a few fragments skip exon 6, which shifts the reading frame, and the 5^*′*^ end of the gene uses a splice donor in exon 1 that differs from the protein-based model (Supplementary Note S4). Neither affects codon 338.

### 2.4 Every *Gyps* species examined carries the stop; no other raptor does

Every vulture assembly available at NCBI on 19 November 2026 was surveyed, together with the reference genome of every other accipitrid species with a long-read-quality assembly, the Peregrine Falcon as an outgroup, and four assemblies added to check particular calls (Methods; Table S9): 45 assemblies of 32 species in all (Table S2). One of them is not independent: the short-read assembly labelled *Accipiter melanochlamys* (GCA_027497775.1) has the same contig N50 (17,712,854 bp), the same number of contigs and the same *CYP2C45* sequence as the Northern Goshawk reference genome (*Astur gentilis*, GCF_929443795.2), and is evidently derived from it; it was kept in the survey but not counted as a species (Supplementary Note S5). In 43 assemblies (30 species) codon 338 could be read. Two could not: the short-read assembly of Rüppell’s Vulture (*G. rueppelli*) contains only four of the nine exons (the codon was read from its raw reads instead; see below), and the alternate haplotype of the White-tailed Eagle lacks the gene (Supplementary Note S5). The codon was read in two independent ways, from the protein-guided gene model and by aligning the *G. africanus* sequence around the codon directly to each locus. The two readings agreed in every assembly except that of the Andean Condor,whose gene model is thrown out of frame by a one-base indel further upstream; there the direct alignment reads CGA (Supplementary Note S5).

TGA was found in all five *Gyps* assemblies. They represent three species and five birds, and come from four laboratories and four sequencing technologies (Nanopore, PacBio, Illumina and DNBseq). It was absent from all 27 non-*Gyps* species (Fig. 4A; Table S1): six other Old World vultures, fifteen eagles, hawks, harriers and allies, five New World vultures and the Peregrine Falcon. Where a species had more than one assembly, they agreed, with one exception that does not involve a stop: the two haplotypes of the same Bearded Vulture read CGA and CAA (Supplementary Note S5). The neighbouring gene *CYP2C23* had an intact reading frame in every assembly, including those of *Gyps*.

### 2.5 The three South Asian species are homozygous for the stop

*G. bengalensis, G. indicus* and *G. tenuirostris* have raw whole-genome reads (roughly 45–52-fold coverage) but no assembly. Assembling a whole genome to read one codon is unnecessary. Instead, the few reads that carry short sequences (18 nucleotides) tiled along exon 7 were pulled out and aligned to the *G. africanus* gene, and the base at codon 338 was read from the alignments (Methods). The same probes, tiled along exon 7 of *CYP2C23*, served as a positive control. The first 60 million read pairs of each run (28–32% of the data) were scanned.

All three species carry TGA, and none shows the intact CGA allele (Table 1; Fig. 4B). For the first position of the codon, the base that differs, 14 independent DNA fragments of *G. bengalensis*, 14 of *G. tenuirostris* and 12 of *G. indicus* read T, and none read C. If an individual carried one intact copy, each fragment would have an even chance of showing it; the chance of seeing no C among 12 to 14 fragments would be 0.5^12^ to 0.5^14^ (2 *×* 10^*−*4^ to 6 *×* 10^*−*5^). Each bird is therefore homozygous for the stop. The control gene was recovered at similar depth (16–33 fragments; Table 1), so the method does find a gene that is present. The probes could not favour the stop allele: every read that covered codon 338 also carried several probes that do not overlap the codon, and would have been recovered without the two probes that do (Table S4).

**Table 1:** Codon 338 of *CYP2C45* read from raw genome reads of the three South Asian *Gyps* species. Fragments are read pairs counted once; bases with quality *≥* 20. The control column counts fragments aligned to exon 7 of the intact neighbouring gene *CYP2C23. P* is the probability of seeing no C at the first codon position among the T-reading fragments if the bird carried one intact copy. For *G. indicus*, a second scan of read pairs 60–110 million added 3 full TGA codons and 4 T reads (Supplementary Note S3); totals are shown.

| Species | Run | Read pairs | Control | Codon 338 |  | Position 1 | *P* |
|---|---|---|---|---|---|---|---|
|  |  | scanned | fragments | TGA | CGA | T : C |  |
| *G. bengalensis* | SRR23198172 | 60 M | 16 | 14 | 0 | 14 : 0 | 6 × 10<sup>−5</sup> |
| *G. tenuirostris* | SRR23198173 | 60 M | 28 | 12 | 0 | 14 : 0 | 6 × 10<sup>−5</sup> |
| *G. indicus* | SRR23198174 | 110 M | 33 | 11 | 0 | 12 : 0 | 2 × 10<sup>−4</sup> |

With these three species and the Cape Vulture transcript, the stop is known in seven of the eight living *Gyps* species, from Africa, Europe, the Himalayas and the Indian subcontinent, and in every case it is the only allele seen. The eighth, Rüppell’s Vulture, is added in the next section.

### 2.6 More birds, and why public population data do not help

Raw genome reads of four further *Gyps* birds were scanned in full with the same probes, filters and counting (Table S18). The assembly of Rüppell’s Vulture misses codon 338, but raw reads of the same museum bird (LACM 61059; 104.8 million read pairs) are public. This bird is homozygous for the stop: all 7 fragments with a complete high-quality codon read TGA and none read CGA (*P* = 0.5^7^ = 0.008 for a heterozygote), and the control gene was recovered as in the other birds (29 fragments). A second Cape Vulture (bird CV01; 82.6 million Ion Torrent reads) and both parents of a captive Griffon Vulture pedigree sequenced to measure mutation rates (Bergeron et al., 2023; 112.8 and 162.3 million read pairs) are also homozygous: 10, 10 and 21 fragments read TGA, none read CGA, and *P ≤* 0.001 in each bird. The stop has now been read in all eight living *Gyps* species and in 13 birds, and the intact allele has not been seen once.

Beyond these birds, public population-scale data could not be used. Two RAD-sequencing studies of Indian vultures are public: 129 *G. indicus, G. bengalensis* and *G. tenuirostris* (BioPro-ject PRJNA1288745) and 15 vultures including *G. bengalensis* and *G. fulvus* (PRJNA1404108). RAD sequencing reads only the DNA next to particular restriction sites. In the six birds tested, from both projects and all three South Asian species, not one read in either read file covered codon 338 (Table S18). The restriction map around the codon explains this. With the enzyme pairs commonly used for double-digest RAD, the codon falls in a fragment that either has the same enzyme at both ends, and so is not sequenced, or is more than 1 kb long, beyond the usual size selection (Table S19). How common the stop is in wild populations therefore still rests on its presence in every bird examined; the PCR–RFLP assay described in the Discussion could measure it directly.

### 2.7 The stop arose on the branch leading to *Gyps*, at least 1.5 million years ago

The stop is present in all eight living *Gyps* species, including *G. bengalensis*, whose lineage was the first to branch off within the genus (Johnson et al., 2006). It is absent from *Necrosyrtes*, the sister genus of *Gyps* (Catanach et al., 2024), *and from every other raptor. The simplest history is therefore a single mutation on the Gyps* stem (Fig. 5A). That branch began when *Gyps* separated from *Necrosyrtes*, 10.6 million years ago in the dated tree of Catanach et al. (2024) (6 million years in the earlier estimate of Mindell et al., 2018), and ended with the common ancestor of the living species, 2.7 (1.5) million years ago. A stop already polymorphic before the split, and later lost by *Necrosyrtes*, cannot be excluded but requires more steps. A finer date from the change of dN/dS along the stem (Meredith et al., 2009) was not possible: the stem carries an estimated five non-synonymous and no synonymous substitutions, so its dN/dS has no upper bound. The stop has thus been present in every *Gyps* lineage for at least 1.5–2.7 million years.

**Figure 5:**
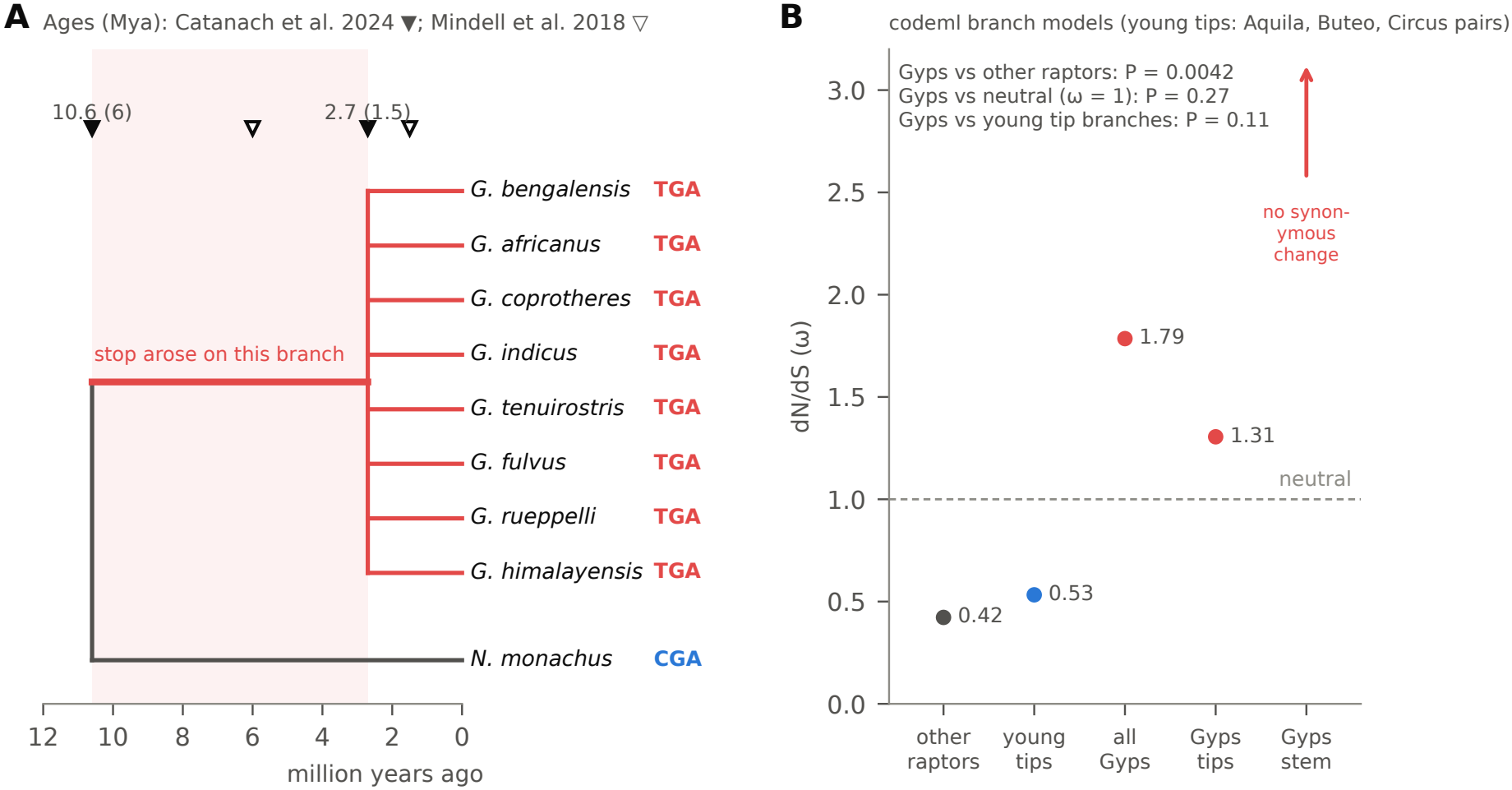
When the stop arose, and how *CYP2C45* has evolved since. **(A)** The stop is present in all eight living *Gyps* species and absent from *Necrosyrtes*, so it arose on the shaded branch. Ages of the *Gyps*–*Necrosyrtes* split and of the *Gyps* crown from Catanach et al. (2024) (filled) and Mindell et al. (2018) (open); relationships within *Gyps* are shown unresolved. **(B)** dN/dS (*ω*) estimated with codeml branch models on the species tree: all non-*Gyps* branches, the tip branches of three young congeneric pairs (control), all *Gyps* branches, the *Gyps* tips and the *Gyps* stem (no synonymous change, so *ω* is unbounded). Dashed line: neutral evolution.

### 2.8 Since the stop, the gene has evolved at about the neutral rate, but the signal is small

If *CYP2C45* lost its function when the stop appeared, amino-acid changes should since have accumulated in *Gyps* about as fast as silent ones (dN/dS, *ω*, near 1), whereas in other raptors purifying selection keeps *ω* well below 1. This was tested on a codon alignment of 27 species, one assembly each, plus the Cape Vulture sequence assembled from its liver RNA-seq (495 codons; Methods; Table S13), with the stop coded as missing.

In branch models on the species tree (Fig. 5B; Table S14), *ω* on the *Gyps* branches was 1.79, against 0.42 on all other branches (likelihood-ratio test, *P* = 0.004), and did not differ from 1 (*P* = 0.27). The four *Gyps* tip branches together had *ω* = 1.31 (not different from 1, *P* = 0.64). There was no sign of positive selection: BUSTED found none on the *Gyps* branches (*P* = 0.25), and aBSREL none on any of the 49 branches of the tree. These numbers fit a gene that no longer has a function. The signal is nevertheless small, because *Gyps* is young: its branches carry an estimated 18 non-synonymous and 4 synonymous substitutions in all, 5% of the length of the tree. RELAX (Wertheim et al., 2015), which lets *ω* vary among sites, gave no consistent answer. Depending on the tree and on whether the stem was included, its relaxation parameter ranged from *k* = 0 (complete relaxation; *Gyps* tips on the species tree, *P* = 0.049) to *k* = 3.1–4.1 (nominally stronger selection; all *Gyps* branches, *P* = 0.003–0.008), and fits of the same model from different random starting values differed by up to 3.9 log-likelihood units (Supplementary Note S7). With some twenty substitutions on the test branches, its three classes of sites cannot be estimated reliably, and RELAX is not interpreted further. Finally, *ω* tends to be raised on any short terminal branch, because slightly harmful changes have not yet been removed (Mugal et al., 2014). On the six tip branches of the three youngest congeneric pairs in the panel (*Aquila, Buteo, Circus*), *ω* was 0.53, no different from the other branches (*P* = 0.66; RELAX *k* = 0.54, *P* = 0.86) but also not significantly lower than in *Gyps* (*P* = 0.11). On the maximum-likelihood gene tree, *ω* on the *Gyps* branches was 1.91 against 0.43 (*P* = 0.035), again not different from 1 (*P* = 0.38) or from the young tip branches (*P* = 0.16) (Table S14). The selection analysis therefore agrees with a loss of function but could not, on its own, establish one; the stop codon and the transcript do that.

## 3 Discussion

The results turn a single report into a genus-wide picture. The stop codon in *CYP2C45* is present in all eight living *Gyps* species and, as far as 13 birds can show, fixed, including the three species whose collapse made diclofenac infamous. It sits in a coding exon, the mature mRNA carries it, and the protein it encodes would lack the residue that binds haem. In practice, every *Gyps* vulture should be treated as having no working *CYP2C45* enzyme.

The view of the earlier reports changes in two ways. The stop does not depend on an unusual retained intron. Intron 7 is spliced as efficiently as the other introns, and the codon lies in exon sequence that all raptors share. Why it was earlier described as intronic is not certain; a gene model built by matching to human *CYP2C19*, which is a different gene, is one possibility. Second, the gene is the bird *CYP2C45*, the most highly expressed of the cytochrome P450 genes measured in chicken liver (Watanabe et al., 2013). The name matters because human *CYP2C19* and *CYP2C9* have different drug preferences; carrying over the substrate list of a particular human enzyme to the bird gene is not justified.

Because the stop is shared by African, European and Asian *Gyps*, the simplest explanation is a single mutation on the branch leading to the genus, between about 11 and 1.5 million years ago, that has since become fixed. Since then the gene has gained amino-acid changes at about the rate expected without selection, as genes without function do, although the genus is too young for this signal to be strong. A gene can be lost without harm when another enzyme covers its job or its job no longer matters. The loss happened millions of years before veterinary drugs existed, so it was not caused by them; whatever *CYP2C45* did, *Gyps* vultures have done without it ever since. What the loss does mean is that the defect is part of what a *Gyps* vulture is. It is not a feature of particular populations, and it cannot be bred out.

The finding fits what is known about diclofenac in *Gyps*: slow elimination and a long half-life (Adawaren et al., 2024; Naidoo and Swan, 2009). It does not prove that *CYP2C45* is the enzyme that clears diclofenac in other birds. That needs an enzyme assay of the intact protein from a tolerant species. The survey also shows clearly that the stop is not the only route to harm. The Steppe Eagle, *Aquila nipalensis*, has died in the wild with diclofenac in its tissues and gout in its organs (Sharma et al., 2014), and so has a Cinereous Vulture, *Aegypius monachus*, in Spain (Herrero-Villar et al., 2021). Both carry an intact *CYP2C45*. Lethal sensitivity therefore occurs without this lesion, as Colston et al. (2025) also argued. The lesion may make *Gyps* more sensitive, or more uniformly so, but other factors, such as dose, other enzymes and the transport of uric acid, must also matter.

For conservation, four practical points follow. First, because the defect is genus-wide and fixed, drug-safety tests on African *Gyps*, already used as stand-ins for the South Asian species (Naidoo et al., 2010; Swan et al., 2006), are genetically justified for this gene: they lack the same enzyme. Second, the lack of the stop in a species is no assurance of safety; the eagle and Cinereous Vulture show otherwise. Third, new veterinary anti-inflammatory drugs that depend on CYP2C-type clearance in mammals deserve particular caution for *Gyps*. Fourth, the stop allele creates a BclI restriction site (TGATCA) that the intact allele (CGATCA) lacks, so birds can be genotyped cheaply by PCR and digestion, for example from shed feathers or the blood samples taken in breeding centres. Primers designed in silico that match every *Gyps* and other Aegypiinae assembly are given in Table S17; they have not been tested at the bench.

### Limitations

Most species are represented by one bird (two for *G. africanus* and *G. coprotheres*, four for *G. fulvus*), and the call for Rüppell’s Vulture rests on seven informative fragments from a museum specimen. The results show that the stop is present and, where raw reads exist, homozygous, and its presence in every bird examined argues that it is fixed; but its frequency in wild populations has not been measured, because the only population-scale data, from RAD sequencing, do not cover the codon. The South Asian genotypes come from about a third of each sequencing run; the remaining reads would add depth but are not needed for the call. The RNA-seq comes from one Cape Vulture and was mapped to a closely related species. The 5^*′*^ end of the gene model is less certain than the rest (Supplementary Note S4), which does not affect exons 3 to 9 or codon 338. Nonsense-mediated decay and the loss of enzyme activity are inferred from well-established rules, not measured. The selection tests have little power, because *Gyps* is young, and the timing rests on published node ages, which differ between studies by nearly a factor of two. Other *CYP2C* -like genes of birds were not examined in vultures. Finally, two non-*Gyps* assemblies show a different stop, at codon 190, on one haplotype only; this could be a real heterozygous variant or an assembly error and is left unresolved (Supplementary Note S6). It does not bear on the *Gyps* result.

## 4 Methods

### Data

All data are public (Tables S9–S10). Genome assemblies were chosen by fixed rules from a snapshot of NCBI Datasets taken on 19 November 2026 (335 assembly records of Accipitridae, Cathartidae, Falconidae, Pandionidae and Sagittariidae): every assembly of every vulture species, after removing duplicate submissions of the same assembly; the NCBI reference genome of each other accipitrid species whose contig N50 is at least 1 Mb; the reference genome of the Peregrine Falcon; and four assemblies added to check specific calls (the second haplotypes of a Hen Harrier and a White-tailed Eagle, and a congener of each). This gave 45 assemblies of 32 species. An assembly whose *CYP2C45* coding sequence and contig N50 were both identical to those of another species’ assembly was treated as a copy of it and not counted (one case; Results). Raw whole-genome reads of *G. bengalensis* (SRR23198172), *G. tenuirostris* (SRR23198173) and *G. indicus* (SRR23198174) are from BioProject PRJNA917070 (National Centre for Biological Sciences; Illumina HiSeq X Ten, 2*×*151 nt). The Cape Vulture liver transcriptome is BioProject PRJNA912371 (University of Pretoria; runs SRR22807161–SRR22807163; HiSeq 2500, 2*×*125 nt). Chicken and human protein sequences and genome annotations were obtained through the NCBI Datasets interface.

### Finding the gene

Twelve chicken P450 proteins, including CYP2C45 (NP_001001752.2) and CYP2C23a (NP_001001616.2), were aligned to each assembly with miniprot 0.18 (Li, 2023) (options -M 3 –outn=5). For each assembly the best *CYP2C45* and *CYP2C23* models were kept, together with 5 kb of flanking sequence. A model was rejected as *CYP2C45* if it fell on the *CYP2C23* locus and matched chicken CYP2C23a better. Coding sequences were joined from the model’s exons and translated without stopping at internal stop codons.

### Reading codon 338

Codon 338 was defined in *G. africanus* as the codon whose first base is position 8,404,731 of contig JBEHDR010000102.1. Each species’ protein was aligned to the *G. africanus* protein (global alignment, BLOSUM62, gap opening *−*11, extension *−*1; Biopython 1.88 (Cock et al., 2009)), and the aligned codon was read from that species’ coding sequence. Because a single-base error upstream can shift a gene model out of frame, the codon was also read directly: the 123-nucleotide *G. africanus* sequence centred on the codon was aligned to each locus on both strands (local alignment; match 2, mismatch *−*3, gap opening *−*5, extension *−*2), and the three aligned bases were read. This frame-independent reading is the one reported; it aligned with 89–100% identity in all 43 assemblies. Splice-site dinucleotides were read from the intron ends of each model.

### RNA-seq

Each read was aligned separately to the 31.8-Mb *G. africanus* contig with minimap2 (Li, 2018) (mappy 2.31, preset splice:sr). Only primary alignments with at least 95% identity over at least 80% of the read were kept, and the two reads of a pair were counted once (fragment counting). For each intron, *S* was the number of fragments spliced exactly across it and *U* the mean number of fragments that ran without a break across its two exon–intron boundaries, with at least 10 nucleotides on each side; intron retention was *U/*(*U* + *S*). Bases at codon 338 were counted when their quality was at least 20.

### Genotyping from raw reads

Fifteen 18-nucleotide probes tiled at 12-nucleotide steps across *CYP2C45* exon 7, and eight across exon 7 of *CYP2C23* as a control, were taken from *G. africanus* (Table S11). Reads containing any probe, on either strand, were extracted from the first 60 million read pairs of each run by exact matching (grep -F). Short, overlapping probes tolerate the small differences between species, and the control locus guards against a silent failure of recruitment. Recruited reads were aligned with minimap2 (preset sr) to 4-kb windows around the two exons; alignments with at least 90% identity over at least 60% of the read were kept. Bases at codon 338 with quality at least 20 were counted once per fragment. Off-target reads, which matched a single probe elsewhere in the genome, failed to align and were discarded. In the analysed portion of each file, read numbers ran without gaps or repeats from 1 to 60,000,000, confirming that the files were intact there (Supplementary Note S3).

### Further birds and RAD-seq data

Raw reads of a second Cape Vulture (SRR12539098; BioProject PRJNA659902; Ion Torrent, single reads) and of the parents of a Griffon Vulture pedigree (SRR17072741, SRR17072739; PRJNA767781; Bergeron et al., 2023), and of the Rüppell’s Vulture whose assembly is in the survey (SRR17999784; PRJNA771872; museum specimen LACM 61059), were obtained from the NCBI Sequence Read Archive with prefetch, because downloads of the ENA copies repeatedly broke off. They were converted with fasterq-dump or, where disk space was short, streamed with fastq-dump in consecutive ranges of spots (SRA Toolkit 3.4.1). Every read was scanned, and the number of reads scanned matched the run records exactly. Reads were recruited, aligned and counted as above. From each of the two RAD-seq projects (PRJNA1288745 and PRJNA1404108), the complete read files of three test birds were scanned in the same way, and reads carrying either 25-nucleotide sequence that flanks codon 338 were counted; all birds of a project were to be scanned if any test bird had such a read. Restriction sites of the enzymes commonly used for RAD sequencing were mapped around the codon on the *G. africanus* contig.

### Gene tree

Thirty-five CYP2 proteins (human CYP2C8, 2C9, 2C18, 2C19, 2E1, 2J2 and 2D6; chicken CYP2C45, 2C18L, 2C23a, 2C23b, 2D49 and 2J23; CYP2C45 and CYP2C23 from eleven raptors) were aligned with FAMSA (Deorowicz et al., 2016). Columns with more than 50% gaps were removed, leaving 494. A maximum-likelihood tree was built with VeryFastTree 4.0.4.1 (Piñeiro et al., 2020), which implements the FastTree-2 method (Price et al., 2010), under the LG model (Le and Gascuel, 2008) with gamma-distributed rates, and SH-like local supports. A neighbour-joining tree (Saitou and Nei, 1987) of Poisson-corrected distances was built with 1,000 bootstrap replicates. Trees were rooted on the CYP2D genes. In *Gyps*, the codon 338 stop was treated as unknown.

### Extended gene tree

Every RefSeq protein annotated as a member of the CYP2C or CYP2H subfamily was taken from the NCBI annotations of thirteen vertebrates (Barn Owl, Tawny Owl, Golden Eagle, Zebra Finch, Mallard, turkey, chicken, American Alligator, Painted Turtle,Green Anole, *Xenopus tropicalis*, mouse and human; the longest protein of each gene; Table S16). Together with the 35 proteins above, and after removing seven exact duplicates, this gave 117 proteins. They were aligned with FAMSA, and columns with more than 50% gaps were removed (494 kept). A maximum-likelihood tree was built with IQ-TREE 3.1.4 (Wong et al., 2026). ModelFinder (Kalyaanamoorthy et al., 2017) chose the model by BIC (JTT+F+R5) among eight protein matrices (LG, WAG, JTT, JTTDCMut, VT, Q.pfam, Q.bird, Q.mammal), with or without empirical frequencies, invariant sites, gamma or FreeRate heterogeneity. Support came from 1,000 ultrafast bootstraps (Hoang et al., 2018) and 1,000 SH-aLRT replicates (Guindon et al., 2010). To test one-to-one orthology directly, the best tree was found under each of four constraints, in which avian CYP2C45 forms a clade with one human gene (CYP2C8, 2C9, 2C18 or 2C19) apart from the other three human genes and the mouse Cyp2c genes. The five trees were compared with the approximately unbiased test (Shimodaira, 2002) (10,000 RELL replicates).

### Selection analysis

One assembly per species was used: the one with the highest contig N50 among those whose *CYP2C45* model was complete (nine exons and a terminal stop), free of frameshifts and of in-frame stops other than the *Gyps* codon 338, and not a copy of another species’ assembly (27 species; Table S13). The Cape Vulture coding sequence was built from the liver RNA-seq: a base was called where at least five fragments covered it and at least 80% agreed (1,293 of 1,488 bases). Proteins were aligned with FAMSA, codons were placed on the protein alignment (495 codons), and stop codons were coded as missing. Two trees were used. The main one is a species tree from published phylogenies, with Accipitridae and Cathartidae relationships as in Catanach et al. (2024), the Peregrine Falcon as outgroup, and relationships within *Gyps* and among *Accipiter, Astur* and *Circus* left unresolved. The second is the maximum-likelihood tree of the codon alignment (IQ-TREE 3.1.4; ModelFinder; 1,000 ultrafast bootstraps). Branch models (Yang, 1998) were fitted with codeml (PAML 4.10.10; Yang, 2007; F3*×*4 codon frequencies; the limit of three descendants per node raised to ten to allow polytomies). The models were: one *ω*; one *ω* for all *Gyps* branches and another for the rest; separate *ω* for the *Gyps* stem and the *Gyps* tips; each with the *Gyps ω* also fixed at 1; and a model with a third *ω* for the tip branches of the three youngest congeneric pairs in the panel (*Aquila chrysaetos* and *A. nipalensis, Buteo buteo* and *B. regalis, Circus aeruginosus* and *C. cyaneus*), as a control for the rise of dN/dS on short branches (Mugal et al., 2014). Nested models were compared by likelihood-ratio tests (*χ*^2^, 1 degree of freedom). HyPhy 2.5.99 (Kosakovsky Pond et al., 2020) was used for RELAX (Wertheim et al., 2015), with three test sets: all *Gyps* branches; the *Gyps* tips with the stem as a nuisance set; or the control tips. HyPhy was also used for BUSTED (Murrell et al., 2015) on the *Gyps* branches and aBSREL (Smith et al., 2015) on all branches.

### Timing

Ages of the *Gyps*–*Necrosyrtes* split and of the *Gyps* crown were taken from the dated phylogeny of Catanach et al. (2024) and, for comparison, from Mindell et al. (2018) as quoted there. The mixed-branch method of Meredith et al. (2009), which places the loss of constraint on a branch from that branch’s *ω*, was applied to the three-ratio model; it needs at least one synonymous substitution on the stem.

### Diagnostic assay (in silico)

Primers flanking codon 338 were designed on the *G. africanus* sequence with primer3 (Untergasser et al., 2012) (product 280–480 bp; *T*_*m*_ 57–63 ^*◦*^C). A pair was kept if it matched every *Gyps* and other Aegypiinae assembly with no mismatch in the five 3^*′*^-terminal bases and at most one elsewhere, and if the two BclI fragments of the stop allele differed clearly in length. Every assembly was then amplified and digested in silico (Table S17).

### Synteny

Gene positions around the chicken (GRCg7b, chromosome 6, 16.3–19.4 Mb) and human (GRCh38, chromosome 10, 94.2–97.0 Mb) *CYP2C* genes were taken from NCBI annotation reports. Fourteen chicken neighbour proteins were aligned to the *G. africanus* contig with miniprot.

## Data and code availability

All sequence data are public at NCBI and ENA under the accessions listed in Tables S9, S10, S16 and S18. The reproducibility archive (code, small data extracts, result tables, figures and L^A^TEX source) is at https://doi.org/10.5281/zenodo.24420380 under CC BY 4.0 (data, text) and MIT (code) licences. Supplementary Note S11 explains how to rerun every analysis from it.

## Acknowledgements

The groups who generated and released the genome and transcriptome data used here are grate-fully acknowledged, in particular the National Centre for Biological Sciences (PRJNA917070), the University of Pretoria (PRJNA912371), the authors of the Indian vulture RAD-seq studies (PRJNA1288745, PRJNA1404108), the groups who released the raw reads of a Cape Vulture (PR-JNA659902), a Griffon Vulture pedigree (PRJNA767781) and Rüppell’s Vulture (PRJNA771872), and the many laboratories and consortia behind the raptor genome assemblies listed in Table S9.

## Use of AI tools

An AI assistant (Claude, Anthropic) was used to help write and run the analysis code and to draft the text. K.S. directed the work and checked the analyses and their outputs. Both authors take full responsibility for the content.

