## Supplementary material for "A stop codon shared by all *Gyps* vultures, including the three Critically Endangered species of South Asia, lies in a coding exon of the bird gene *CYP2C45*": Supplimentary material

Krishnendu Sinha

**Supplementary Figures** **6**

**Supplementary Tables** **8**

All tables are also provided, unabridged, in `Supplementary_Tables_S1-S12.xlsx`. Every number below is produced by the scripts in the reproducibility archive (`https://doi.org/10.5281/zenodo.24420380`); Note S7 explains how to rerun them.

#### Supplementary Notes

##### Note S1. The stop codon is the variant reported earlier

Adawaren et al. (2024) reported a premature stop codon in the Cape vulture gene they called *CYP2C19* and described it as lying in intron 7 and retained in the mature mRNA. Colston et al. (2025) gave a 66-nucleotide Cape vulture sequence containing the stop and listed where it occurs in three other *Gyps* assemblies. Each of their coordinates contains the codon studied here, 36 nucleotides from the start of their sequence:

| Species | Contig | 66-nt match (Colston et al., 2025) | First base of TGA (this study) |
|---|---|---|---|
| *G. africanus* | JBEHDR010000102.1 | 8,404,695–8,404,760 (+) | 8,404,731 |
| *G. fulvus* | JAOEID010000099.1 | 20,115,051–20,115,116 (+) | 20,115,087 |
| *G. himalayensis* | JACFZI010000879.1 | 709,499–709,434 (−) | 709,463 |

The published sequence begins with the last base of intron 6 (the G of the AG acceptor at 8,404,694–8,404,695) and continues with the first 65 bases of exon 7. The TGA occupies bases 36–38 of exon 7. The variant is therefore the same; only its description differs. Here it lies in coding exon 7, and intron 7, which starts 150 bases further downstream, is efficiently spliced (main text, Fig. 3).

#### Note S2. Why the gene is called *CYP2C45*

Bird *CYP2C* genes arose by duplications separate from those that produced the human genes (Watanabe et al., 2013). In chicken, the gene that Watanabe et al. (2013) called *CYP2C45* is RefSeq NP_001001752.2 (NCBI Gene 414833, current symbol *CYP2C18*, synonym *CYP2C45*). A second copy, 99.4% identical at the protein level, sits next to it, 9 kb upstream (start to start) (*CYP2C18L*, Gene 100858007, XP_015144012.2); this duplication is specific to the chicken lineage or to its assembly, because each raptor assembly has a single copy. The second bird gene, *CYP2C23* (*CYP2C23a/b* in chicken, formerly *CYP2H1*/*CYP2H2*), lies about 1 Mb away near *ABCG2* and *ALOX5*.

The vulture protein matches chicken CYP2C45 far better than CYP2C23a (81.0% versus 55.6% identity in global pairwise alignment; 80.8% to CYP2C45 in the miniprot alignment), lies at the same position between *ZP4* and *TLL2* (Table S6 in the workbook), and groups with chicken CYP2C45 in both trees (Fig. 1A; Fig. S1; Table S7). The four human *CYP2C* genes occupy the same neighbourhood in humans and form their own clade, so none of them is a one-to-one counterpart of the bird gene. In the maximum-likelihood tree, human CYP2E1 falls next to the human CYP2C group rather than outside all CYP2C genes; this placement is weakly determined and has no bearing on the conclusions. Codon 338 aligns with a conserved arginine, residue 333 in human CYP2C8, CYP2C9 and CYP2C19 (Table S8).

#### Note S3. Reading genotypes from raw reads

**Probe design.** Fifteen overlapping 18-nucleotide probes (12-nucleotide steps) cover exon 7 of *CYP2C45*; short, overlapping probes still match when a few bases differ between species. The same design across exon 7 of *CYP2C23* acts as a positive control: if the control is recovered and the target is not, the target is missing; if neither is recovered, the method has failed (Table S11). The probes are generated from the *G. africanus* reference by `scripts/05d_make_probes.py`.

**Which reads count.** About a third to two thirds of recruited reads matched a single probe that also occurs elsewhere in the genome; they did not align to either gene window and were discarded. Reads that did align were counted per fragment (both reads of a pair together), and only bases with quality ≥20 were used (Table S3).

**Allele bias.** Two of the fifteen exon-7 probes overlap codon 338 and carry TGA, so a CGA read would not match them. But every read covering the codon also carried other probes, and would have been recovered without those two (Table S4). The recruitment cannot hide an intact allele.

**File integrity.** The downloaded FASTQ files are 6–7% larger than the byte counts recorded by ENA. The reason was not resolved (a different compression or a resumed download are possible). Instead, the part that was analysed was checked. In each of the six files, the first 60,000,000 records have read numbers that rise strictly from 1 to 60,000,000, with no repeats or gaps, and no read name occurs twice among the recruited reads of any file. The analysed portion is therefore intact (`scripts/05c_check_fastq_integrity.sh`). The base calls do not depend on depth, and the counts are counts of distinct fragments.

**A follow-up scan of *G. indicus*.** In the first scan, no *G. indicus* fragment crossed the 3′ end of exon 7, whereas 6–9 did in the other species, and two short fragments (inserts of about 60–75 bp, with adapter sequence in the reads) aligned only partly. To test whether *G. indicus* has a structural change there, read pairs 60–110 million were scanned with the same probes plus probes built from the unaligned parts of those two fragments. Nine fragments now crossed the end of exon 7 normally, the reads matching the new probes were off-target, and three more fragments read TGA at codon 338. The earlier gap was chance, and the two odd fragments were library chimeras. The follow-up reads are included in the archive and in Table S3; the four extra probes, and the reads they were taken from, are listed in `data/probes/indicus_followup_extra_kmers.tsv`.

#### Note S4. Details of the RNA-seq analysis

Of 8,645,827 read pairs, 1,217 aligned to the two genes (89, 563 and 565 in runs SRR22807161–3) after the filters (primary alignment, ≥95% identity over ≥80% of the read). Splicing of every intron is in Table S5. Three further observations:

- **The 5′ end differs from the protein-based model.** No fragment splices across the predicted intron 1 (8,401,420–8,401,760). Instead, four fragments join a donor inside predicted exon 1 (after 8,401,369) to an acceptor at 8,401,456, and coverage continues into part of the predicted intron. Coverage here is low (2.6 fragments per base), so the first exon(s) of the vulture transcript cannot be defined precisely. Codon numbers in this paper refer to the 496-codon reading frame built from the chicken protein; exons 3–9, codon 338 and all conclusions are unaffected.
- **A minor isoform skips exon 6.** Four fragments join exon 5 directly to exon 7, against 7–8 for the normal junctions. Exon 6 is 142 nucleotides long, so skipping it shifts the reading frame. This isoform therefore cannot restore a working protein.
- **Splicing beyond the coding region.** Twenty fragments are spliced between 8,408,890 and 8,409,330, 1.8 kb beyond the stop codon of the reading frame, with a few further junctions downstream. These may belong to an untranslated 3′ region or to another transcript; they were not analysed.

Both genes show more coverage toward their 3′ ends (Fig. 3C; Fig. S2), a common feature of poly(A)-selected libraries. The expression comparison uses the mean over all exons of each gene, so it is affected similarly in both.

#### Note S5. The assembly panel, and assemblies that need comment

The panel was chosen by fixed rules (main text, Methods) from the NCBI snapshot of 19 November 2026, which is included in the archive together with the script that applies the rules (`scripts/00_select_assembli` Table S9 gives the rule that admitted each assembly. Short-read assemblies of non-vulture accipitrids (more than 100 species, most with contig N50 below 50 kb) were not included, because fragmented gene models are hard to read reliably; vultures were included whatever the quality of their assemblies.

- ***Gyps rueppelli*** (GCA_027575145.1). This short-read assembly has a contig N50 of 2.2 kb; only four exons (197 codons) of *CYP2C45* were recovered and codon 338 falls in a gap. Rüppell's Vulture is the only *Gyps* species not genotyped.
- ***Haliaeetus albicilla* alternate haplotype** (GCA_947461935.1, 693 Mb, about half a genome). The best match to chicken CYP2C45 was the *CYP2C23* locus (56.5% identity,

against 76.9% to CYP2C23a), so *CYP2C45* is simply not in this partial assembly. The pipeline rejects such cases automatically.

- ***Vultur gryphus*** (GCA_039700855.2). The gene model contains a one-base indel in exon 3, which shifts the reading frame and produces many apparent stop codons downstream; the protein-based reading of codon 338 is therefore wrong (CTG). The direct nucleotide alignment reads CGA. A single-base indel in a short-read assembly (contig N50 22 kb) is most likely an assembly error.

- ***Gypaetus barbatus***. The primary (GCA_028022735.1) and alternate (GCA_028017155.1) haplotypes of the same bird read CGA and CAA at codon 338. Either the bird is heterozygous or one haplotype carries a base error; neither allele is a stop.

- ***Cathartes aura*** (GCA_000699945.1). This 2014 short-read assembly has ten internal stop codons in codons 45–146 of *CYP2C45*, consistent with assembly errors in a fragmented region; codon 338 (CGA) lies in clean sequence and both readings agree.

- ***Sarcogyps calvus*** GCA_040893125.1 is 2.9 Gb long (probably both haplotypes together) and too large to index in 7 GB of memory; it was aligned in three parts.

- ***Falco peregrinus*** carries CAA (glutamine) at codon 338.

- ***Gyps fulvus*** and ***G. africanus*** are each represented by two assemblies of different birds (different BioSamples); all four read TGA.

#### Note S6. A second stop, at codon 190, in two non-*Gyps* assemblies

The primary assemblies of the Hen Harrier (*Circus cyaneus*, GCA_987196875.1) and the White-tailed Eagle (*Haliaeetus albicilla*, GCF_947461875.1) carry TGA at codon 190 of *CYP2C45*, another CpG arginine codon. The second haplotype of the same Hen Harrier (GCA_987196495.1) reads CGA, and the congeners *C. aeruginosus* and *H. leucocephalus* read CGA; the alternate White-tailed Eagle haplotype lacks the gene (Note S5). A difference between the two haplotypes of one bird fits either a real heterozygous loss-of-function allele or a base error in one haplotype, and the assemblies alone cannot tell these apart; raw reads of these individuals would. Neither species has been tested with diclofenac. These calls do not affect codon 338 or the *Gyps* result and are reported for completeness.

#### Note S7. Reproducing the analysis

The archive contains numbered scripts, the small data they need, and a record of every output. On a Linux computer with Python 3.10 or later (or in the container described by the archive's `Dockerfile`):

```
pip install -r requirements.txt
bash run_all.sh quick     # ~2 minutes, offline: all numbers, tables, figures and both PDFs
bash run_all.sh full      # 1-2 hours: first rebuilds every input from public data
bash scripts/run_wgs.sh /path/to/wgs_reads [download]   # the South Asian read extracts
```

Quick mode starts from the gene models and locus sequences saved by the survey, the RNA-seq read pairs that map to the two genes, and the reads recruited from the three South Asian genomes. Full mode first re-downloads the reference proteins and annotations and compares them with the archived copies, re-applies the selection rules to the NCBI snapshot, re-surveys the 45 assemblies and re-maps all RNA-seq reads (MD5-checked downloads; needs `git`, `make` and a C compiler to build miniprot). `run_wgs.sh` checks the integrity of each raw FASTQ file, re-extracts the reads (including the *G. indicus* follow-up scan) and reports whether they are identical to the archived extracts. Both modes end with `scripts/11_check_results.py`, which compares every reported number with the paper and the SHA-256 checksum of every output file with the archived record. With the software versions given in `requirements.txt` and TeX Live 2023, every output, including the figures and both PDFs, is byte-identical to the archived files.

#### Supplementary Figures

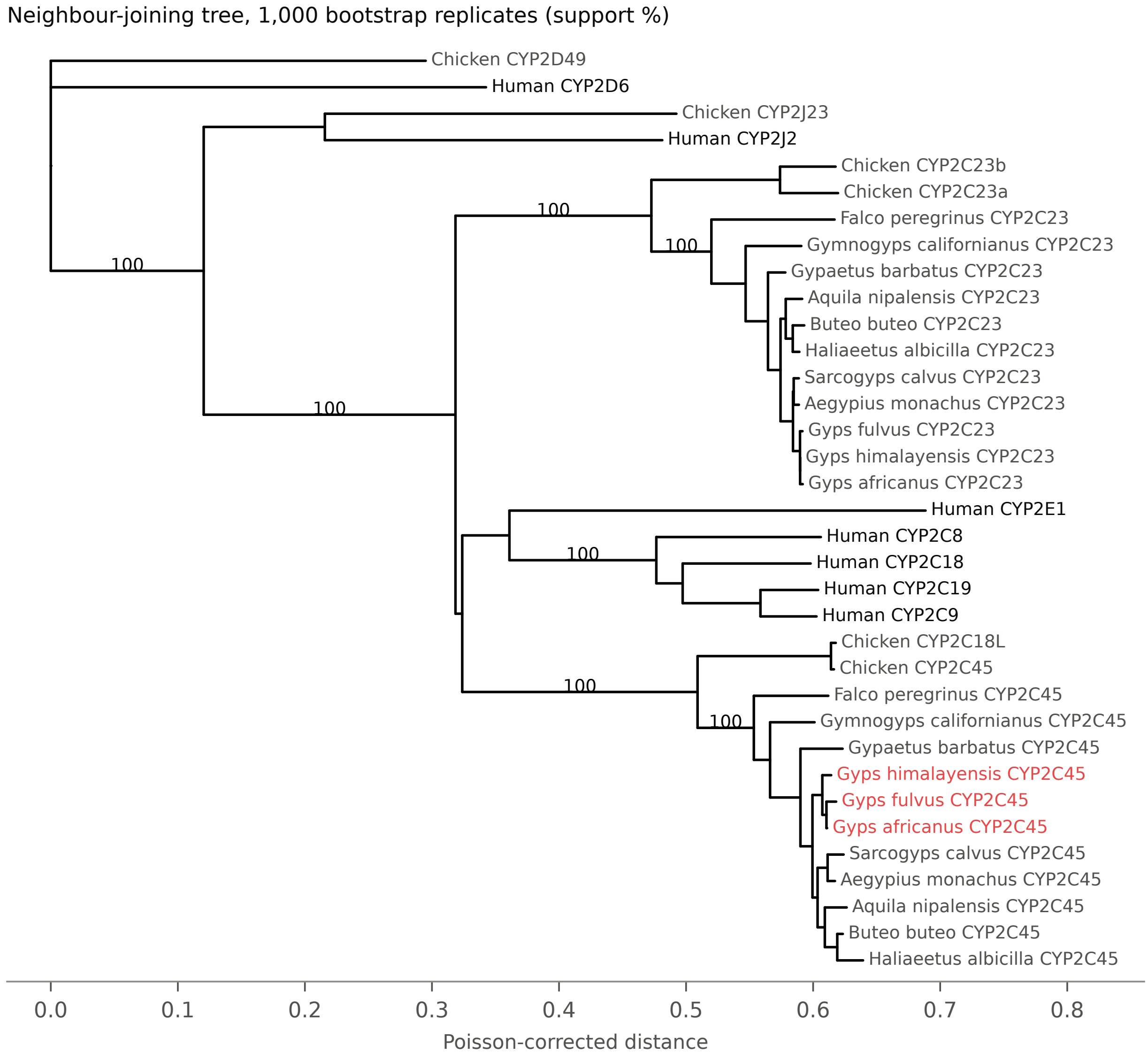

**Figure S1: Neighbour-joining tree of the same 35 CYP2 proteins as Fig. 1A.** Poisson-corrected distances, 494 alignment columns, 1,000 bootstrap replicates; support shown for main branches. *Gyps* CYP2C45 in red.

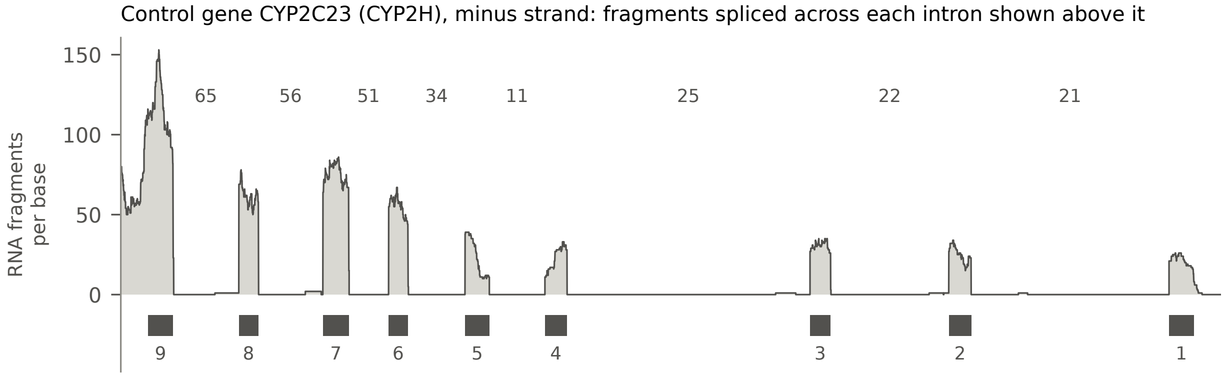

**Figure S2: RNA-seq coverage of the control gene *CYP2C23*** in the Cape vulture liver, with the number of fragments spliced across each intron. Exons are numbered 5′ to 3′ along the gene, which is on the minus strand.

### Supplementary Tables

Tables S6 (synteny coordinates), S9 (assembly list) and S12 (diclofenac evidence) are in the workbook only.

**Table S1: Codon 338 of *CYP2C45* in 33 species.** One row per species; all assemblies in Table S2.

| Group | Species | Codon | Residue | Data | Diclofenac (reference) |
|---|---|---|---|---|---|
| *Gyps* | *Gyps africanus* | TGA | stop | assembly GCA_041146555.1; assembly GCA_041146915.1 | susceptible (Swan et al. 2006a) |
| *Gyps* | *Gyps bengalensis* | TGA | stop | WGS reads SRR23198172 | susceptible (Oaks et al. 2004) |
| *Gyps* | *Gyps coprotheres* | TGA | stop | liver RNA-seq PRJNA912371 | susceptible (Naidoo et al. 2009) |
| *Gyps* | *Gyps fulvus* | TGA | stop | assembly GCA_025727715.1; assembly GCA_056578145.1 | susceptible (Swan et al. 2006a) |
| *Gyps* | *Gyps himalayensis* | TGA | stop | assembly GCA_021398385.1 | untested |
| *Gyps* | *Gyps indicus* | TGA | stop | WGS reads SRR23198174 | susceptible (Shultz et al. 2004) |
| *Gyps* | *Gyps tenuirostris* | TGA | stop | WGS reads SRR23198173 | susceptible (Swan et al. 2006a; Green et al. 2004) |
| Other OW vulture | *Aegypius monachus* | CGA | Arg | assembly GCA_025309335.1; assembly GCA_045787555.1; assembly GCA_982185325.1 | susceptible (Herrero-Villar et al. 2021) |
| Other OW vulture | *Gypaetus barbatus* | CGA/CAA | Arg/Gln | assembly GCA_028017155.1; assembly GCA_028022735.1 | untested |
| Other OW vulture | *Gypohierax angolensis* | CGA | Arg | assembly GCA_055772255.1 | untested |
| Other OW vulture | *Necrosyrtes monachus* | CGA | Arg | assembly GCA_037953595.1; assembly GCA_037953605.1 | untested |
| Other OW vulture | *Neophron percnopterus* | CGA | Arg | assembly GCA_045787405.1; assembly GCA_982185345.1 | untested |
| Other OW vulture | *Sarcogyps calvus* | CGA | Arg | assembly GCA_040869245.1; assembly GCA_040893125.1; assembly GCA_055772215.1 | untested |
| Other accipitrid | *Accipiter melanochlamys* | CGA | Arg | assembly GCA_027497775.1 | untested |
| Other accipitrid | *Accipiter nisus* | CGA | Arg | assembly GCA_978019815.1 | untested |
| Other accipitrid | *Aquila chrysaetos* | CGA | Arg | assembly GCF_900496995.4 | untested |
| Other accipitrid | *Aquila nipalensis* | CGA | Arg | assembly GCA_058581595.1 | susceptible (Sharma et al. 2014) |
| Other accipitrid | *Astur gentilis* | CGA | Arg | assembly GCF_929443795.2 | untested |
| Other accipitrid | *Buteo buteo* | CGA | Arg | assembly GCF_964188355.1 | untested |
| Other accipitrid | *Buteo regalis* | CGA | Arg | assembly GCA_037951015.2 | untested |
| Other accipitrid | *Circus aeruginosus* | CGA | Arg | assembly GCA_982553335.1 | untested |
| Other accipitrid | *Circus cyaneus* | CGA | Arg | assembly GCA_987196495.1; assembly GCA_987196875.1 | untested |
| Other accipitrid | *Haliaeetus albicilla* | CGA | Arg | assembly GCF_947461875.1 | untested |

| Group | Species | Codon | Residue | Data | Diclofenac (reference) |
|---|---|---|---|---|---|
| Other accipitrid | *Haliaeetus leucocephalus* | CGA | Arg | assembly GCF_000737465.1 | untested |
| Other accipitrid | *Harpia harpyja* | CGA | Arg | assembly GCF_026419915.1 | untested |
| Other accipitrid | *Morphnus guianensis* | CGA | Arg | assembly GCA_045345515.1 | untested |
| Other accipitrid | *Pithecophaga jefferyi* | CGA | Arg | assembly GCA_054553165.1 | untested |
| Other accipitrid | *Polyboroides typus* | CGA | Arg | assembly GCA_060712505.1 | untested |
| Other accipitrid | *Spilornis perplexus* | CGA | Arg | assembly GCA_054655575.1 | untested |
| NW vulture | *Cathartes aura* | CGA | Arg | assembly GCA_000699945.1 | tolerant (Rattner et al. 2008) |
| NW vulture | *Coragyps atratus* | CGA | Arg | assembly GCA_055772195.1 | untested |
| NW vulture | *Gymnogyps californianus* | CGA | Arg | assembly GCF_018139145.2 | untested |
| NW vulture | *Sarcoramphus papa* | CGA | Arg | assembly GCA_027580115.1; assembly GCA_037950955.1; assembly GCA_037962945.1 | untested |
| NW vulture | *Vultur gryphus* | CGA | Arg | assembly GCA_039700855.2 | untested |
| Falcon | *Falco peregrinus* | CAA | Gln | assembly GCF_023634155.1 | untested |

**Table S2: *CYP2C45* and *CYP2C23* reconstructed from 34 assemblies.** Codon/aa: codon 338 and its amino acid (* = stop). Sites: donor–acceptor dinucleotides of introns 6 and 7. Internal stops are listed by codon number. Full table with coordinates, identities, sequence context and exon lengths: Supplementary_Tables_S1-S12.xlsx, sheet S2.

| Assembly | Accession | Codon | aa | Exons | Codons | Intron 6/7 sites | Internal stops (CYP2C45) | Stops CYP2C23 |
|---|---|---|---|---|---|---|---|---|
| Gyps_africanus | GCA_041146555.1 | TGA | * | 9 | 496 | GT-AG/GT-AG | 338 | none |
| Gyps_africanus_2 | GCA_041146915.1 | TGA | * | 9 | 496 | GT-AG/GT-AG | 338 | none |
| Gyps_fulvus | GCA_025727715.1 | TGA | * | 9 | 496 | GT-AG/GT-AG | 338 | none |
| Gyps_fulvus_2 | GCA_056578145.1 | TGA | * | 9 | 496 | GT-AG/GT-AG | 338 | none |
| Gyps_himalayensis | GCA_021398385.1 | TGA | * | 9 | 496 | GT-AG/GT-AG | 338 | none |
| Gyps_rueppelli | GCA_027575145.1 | gap | - | 4 | 197 | – | none | none |
| Aegypius_monachus_2 | GCA_025309335.1 | CGA | R | 9 | 496 | GT-AG/GT-AG | none | none |
| Aegypius_monachus_3 | GCA_045787555.1 | CGA | R | 9 | 496 | GT-AG/GT-AG | none | none |
| Aegypius_monachus | GCA_982185325.1 | CGA | R | 9 | 496 | GT-AG/GT-AG | none | none |
| Gypaetus_barbatus_2 | GCA_028017155.1 | CAA | Q | 9 | 495 | GT-AG/GT-AG | none | none |
| Gypaetus_barbatus | GCA_028022735.1 | CGA | R | 9 | 495 | GT-AG/GT-AG | none | none |
| Gypohierax_angolensis | GCA_055772255.1 | CGA | R | 9 | 495 | GT-AG/GT-AG | none | none |
| Necrosyrtes_monachus | GCA_037953595.1 | CGA | R | 9 | 496 | GT-AG/GT-AG | none | none |
| Necrosyrtes_monachus_2 | GCA_037953605.1 | CGA | R | 9 | 496 | GT-AG/GT-AG | none | none |
| Neophron_percnopterus_2 | GCA_045787405.1 | CGA | R | 8 | 417 | GT-AG/GT-AG | none | none |
| Neophron_percnopterus | GCA_982185345.1 | CGA | R | 9 | 495 | GT-AG/GT-AG | none | none |
| Sarcogyps_calvus | GCA_040869245.1 | CGA | R | 9 | 496 | GT-AG/GT-AG | none | none |
| Sarcogyps_calvus_2 | GCA_040893125.1 | CGA | R | 9 | 496 | GT-AG/GT-AG | none | none |
| Sarcogyps_calvus_3 | GCA_055772215.1 | CGA | R | 9 | 496 | GT-AG/GT-AG | none | none |
| Accipiter_melanochlamys | GCA_027497775.1 | CGA | R | 9 | 495 | GT-AG/GT-AG | none | none |
| Accipiter_nisus | GCA_978019815.1 | CGA | R | 9 | 495 | GT-AG/GT-AG | none | none |
| Aquila_chrysaetos | GCF_900496995.4 | CGA | R | 9 | 495 | GT-AG/GT-AG | none | none |
| Aquila_nipalensis | GCA_058581595.1 | CGA | R | 9 | 495 | GT-AG/GT-AG | none | none |
| Astur_gentilis | GCF_929443795.2 | CGA | R | 9 | 495 | GT-AG/GT-AG | none | none |
| Buteo_buteo | GCF_964188355.1 | CGA | R | 9 | 495 | GT-AG/GT-AG | none | none |
| Buteo_regalis | GCA_037951015.2 | CGA | R | 9 | 495 | GT-AG/GT-AG | none | none |
| Circus_aeruginosus | GCA_982553335.1 | CGA | R | 9 | 495 | GT-AG/GT-AG | none | none |

| Assembly | Accession | Codon | aa | Exons | Codons | Intron 6/7 sites | Internal stops (CYP2C45) | Stops CYP2C23 |
|---|---|---|---|---|---|---|---|---|
| Circus_cyaneus_hap2 | GCA_987196495.1 | CGA | R | 9 | 495 | GT-AG/GT-AG | none | none |
| Circus_cyaneus | GCA_987196875.1 | CGA | R | 9 | 495 | GT-AG/GT-AG | 190 | none |
| Haliaeetus_albicilla_alt | GCA_947461935.1 | CYP2C45 absent from assembly (best hit is the CYP2C23 locus, identity 0.5648) | | | | | | |
| Haliaeetus_albicilla | GCF_947461875.1 | CGA | R | 9 | 495 | GT-AG/GT-AG | 190 | none |
| Haliaeetus_leucocephalus | GCF_000737465.1 | CGA | R | 9 | 495 | GT-AG/GT-AG | none | none |
| Harpia_harpyja | GCF_026419915.1 | CGA | R | 9 | 495 | GT-AG/GT-AG | none | none |
| Morphnus_guianensis | GCA_045345515.1 | CGA | R | 9 | 495 | GT-AG/GT-AG | none | none |
| Pithecophaga_jefferyi | GCA_054553165.1 | CGA | R | 9 | 495 | GT-AG/GT-AG | none | none |
| Polyboroides_typus | GCA_060712505.1 | CGA | R | 9 | 495 | GT-AG/GT-AG | none | none |
| Spilornis_perplexus | GCA_054655575.1 | CGA | R | 9 | 495 | GT-AG/GT-AG | none | none |
| Cathartes_aura | GCA_000699945.1 | CGA | R | 9 | 495 | GT-AG/GT-AG | 45, 56, 63, 76, 85, 93, 94, 111, 119, 146 | none |
| Coragyps_atratus | GCA_055772195.1 | CGA | R | 9 | 495 | GT-AG/GT-AG | none | none |
| Gymnogyps_californianus | GCF_018139145.2 | CGA | R | 9 | 495 | GT-AG/GT-AG | none | none |
| Sarcoramphus_papa_2 | GCA_027580115.1 | CGA | R | 9 | 495 | GT-AG/GT-AG | none | none |
| Sarcoramphus_papa_3 | GCA_037950955.1 | CGA | R | 9 | 495 | GT-AG/GT-AG | none | none |
| Sarcoramphus_papa | GCA_037962945.1 | CGA | R | 9 | 495 | GT-AG/GT-AG | none | none |
| Vultur_gryphus | GCA_039700855.2 | CGA | R | 9 | 494 | GT-AG/GT-AG | 174, 201, 202, 205, 211, 212, 244, 252, 259, 288, 291, 292, 310, 334, 374, 417, 462, 473, 477 | none |
| Falco_peregrinus | GCF_023634155.1 | CAA | Q | 9 | 495 | GT-AG/GT-AG | none | none |

**Table S3: Read recruitment and genotypes in the three South Asian species.** Frags: read p counted once. Full codon: fragments with all three bases of codon 338 at quality $\geq$20. P: probability o CGA among the full-codon fragments if the bird were heterozygous.

| Species | Run | Pairs | Recruited reads | CYP2C45 frags | CYP2C23 frags | Frags over exon 7 end | Full codon | TGA |
|---|---|---|---|---|---|---|---|---|
| *Gyps bengalensis* | SRR23198172 | 1-60 M | 75 | 27 | 16 | 9 | 14 | 14 |
| *Gyps tenuirostris* | SRR23198173 | 1-60 M | 88 | 27 | 28 | 6 | 12 | 12 |
| *Gyps indicus* | SRR23198174 | 1-60 M | 75 | 9 | 12 | 0 | 8 | 8 |
| *Gyps indicus (follow-up)* | SRR23198174 | 60-110 M | 88 | 14 | 21 | 9 | 3 | 3 |

**Table S4: The probes cannot favour the stop allele.** Every read that covers codon 338 also carries at least one probe that does not overlap the codon.

| Run | Read pairs | Reads covering codon 338 | Recovered without the two codon probes |
|---|---|---|---|
| SRR23198172 | 1-60 M | 16 | 16 |
| SRR23198173 | 1-60 M | 16 | 16 |
| SRR23198174 | 1-60 M | 8 | 8 |
| SRR23198174 | 60-110 M | 4 | 4 |

**Table S5: Splicing of every intron in the Cape vulture liver.** Coordinates on *G. africanus* contig JBEHDR010000102.1. S: fragments spliced exactly across the intron. U: fragments running unbroken across the left/right exon–intron boundary. IR = $\bar{U}/(\bar{U}+S)$. Introns are numbered 5′ to 3′ along each gene (CYP2C23 is on the minus strand).

| Gene | Intron | Start | End | Length | S | U left | U right | IR | Intron depth |
|---|---|---|---|---|---|---|---|---|---|
| CYP2C45 | 1 | 8401420 | 8401760 | 341 | 0 | 0 | 2 | 1.0 | 2.78 |
| CYP2C45 | 2 | 8401924 | 8402414 | 491 | 4 | 3 | 2 | 0.385 | 0.97 |
| CYP2C45 | 3 | 8402565 | 8402866 | 302 | 10 | 0 | 1 | 0.048 | 0.25 |
| CYP2C45 | 4 | 8403028 | 8403420 | 393 | 11 | 1 | 0 | 0.043 | 0.2 |
| CYP2C45 | 5 | 8403601 | 8404124 | 524 | 7 | 0 | 0 | 0.0 | 0.01 |
| CYP2C45 | 6 | 8404267 | 8404695 | 429 | 8 | 0 | 0 | 0.0 | 0.7 |

| Gene | Intron | Start | End | Length | S | U left | U right | IR | Intron depth |
|---|---|---|---|---|---|---|---|---|---|
| CYP2C45 | 7 | 8404884 | 8406089 | 1206 | 24 | 0 | 5 | 0.094 | 1.0 |
| CYP2C45 | 8 | 8406232 | 8406885 | 654 | 32 | 4 | 1 | 0.072 | 1.18 |
| CYP2C23 | 1 | 9677859 | 9679303 | 1445 | 21 | 0 | 0 | 0.0 | 0.05 |
| CYP2C23 | 2 | 9676832 | 9677695 | 864 | 22 | 0 | 1 | 0.022 | 0.19 |
| CYP2C23 | 3 | 9674907 | 9676681 | 1775 | 25 | 0 | 0 | 0.0 | 0.08 |
| CYP2C23 | 4 | 9674340 | 9674745 | 406 | 11 | 0 | 0 | 0.0 | 0 |
| CYP2C23 | 5 | 9673746 | 9674162 | 417 | 34 | 0 | 0 | 0.0 | 0.05 |
| CYP2C23 | 6 | 9673313 | 9673603 | 291 | 51 | 0 | 0 | 0.0 | 0.16 |
| CYP2C23 | 7 | 9672653 | 9673124 | 472 | 56 | 0 | 0 | 0.0 | 0.53 |
| CYP2C23 | 8 | 9672030 | 9672510 | 481 | 65 | 0 | 1 | 0.008 | 0.58 |

**Table S7: Support for key groupings in the CYP2 protein trees.**

| Grouping tested | NJ bootstrap (%) | ML SH-like support (%) |
|---|---|---|
| avian CYP2C45 clade | 100 | 88 |
| avian CYP2C23 (CYP2H) clade | 100 | 100 |
| human CYP2C clade | 100 | 100 |
| avian CYP2C45 + human CYP2C19 only | 0 | not in tree |
| avian CYP2C45 + avian CYP2C23 | 0 | not in tree |
| all CYP2C (avian + human) | 0 | not in tree |

**Table S8: Position of codon 338 in chicken and human proteins.** The vulture haem-binding cysteine is residue 440.

| Protein | Residue aligned to vulture codon 338 | Amino acid | Haem-binding Cys |
|---|---|---|---|
| chicken CYP2C45 (NP_001001752.2) | 337 | R | 439 |
| human CYP2C19 (NP_000760.1) | 333 | R | 435 |
| human CYP2C9 (NP_000762.2) | 333 | R | 435 |
| human CYP2C8 (NP_000761.3) | 333 | R | 435 |

**Table S10: Read data used.** MD5 checksums of each FASTQ file are in config/runs.tsv of the archive.

| Run | Type | Species | BioProject | Instrument | Read pairs | Pairs analysed |
|---|---|---|---|---|---|---|
| SRR23198172 | WGS | *Gyps bengalensis* | PRJNA917070 | HiSeq X Ten | 209,357,369 | 60,000,000 |
| SRR23198173 | WGS | *Gyps tenuirostris* | PRJNA917070 | HiSeq X Ten | 210,186,774 | 60,000,000 |
| SRR23198174 | WGS | *Gyps indicus* | PRJNA917070 | HiSeq X Ten | 186,056,745 | 60,000,000 |
| SRR22807161 | RNA-Seq | *Gyps coprotheres* | PRJNA912371 | Illumina HiSeq 2500 | 645,827 | 645,827 |
| SRR22807162 | RNA-Seq | *Gyps coprotheres* | PRJNA912371 | Illumina HiSeq 2500 | 4,000,000 | 4,000,000 |
| SRR22807163 | RNA-Seq | *Gyps coprotheres* | PRJNA912371 | Illumina HiSeq 2500 | 4,000,000 | 4,000,000 |

**Table S11: Probes used to recruit reads.** Each probe and its reverse complement were searched (46 strings).

| Set | Probe (5′ →3′) | Start on JBEHDR010000102.1 |
|---|---|---|
| CYP2C45_exon7 | `AGAAAGTTCAAGAAGAGA` | 8404696 |
| CYP2C45_exon7 | `AAGAGATTGACCAGGTAG` | 8404708 |
| CYP2C45_exon7 | `AGGTAGTGGGATGATCAC` | 8404720 |
| CYP2C45_exon7 | `GATCACGAAGACCTTGTG` | 8404732 |
| CYP2C45_exon7 | `CTTGTGTGGCTCATCAGA` | 8404744 |

| Set | Probe (5′ →3′) | Start on JBEHDR010000102.1 |
|---|---|---|
| CYP2C45_exon7 | ATCAGACCCAGATGCCCT | 8404756 |
| CYP2C45_exon7 | TGCCCTACACAGATGCAG | 8404768 |
| CYP2C45_exon7 | ATGCAGTGGTCCATGAAA | 8404780 |
| CYP2C45_exon7 | ATGAAATCCAGCGCTTCA | 8404792 |
| CYP2C45_exon7 | GCTTCATCTCCCTCATCC | 8404804 |
| CYP2C45_exon7 | TCATCCCCCTGGGTCTCC | 8404816 |
| CYP2C45_exon7 | GTCTCCCTCACACTGTGA | 8404828 |
| CYP2C45_exon7 | CTGTGACCAAAGACACCG | 8404840 |
| CYP2C45_exon7 | ACACCGGCTTCAGAGAGT | 8404852 |
| CYP2C45_exon7 | GAGAGTATGTCATTCCCA | 8404864 |
| CYP2C23_control | CTTGGGGATAAAATAGTC | 9673125 |
| CYP2C23_control | CTTGATGTCTTTGATCAC | 9673149 |
| CYP2C23_control | TGGGACATTAAGTGGAAG | 9673173 |
| CYP2C23_control | GATGAATCTCTGGATTTC | 9673197 |
| CYP2C23_control | CACAGCATCTGTGTAGGG | 9673221 |
| CYP2C23_control | GCTCCGATCTGCCATGCA | 9673245 |
| CYP2C23_control | TCGGTCTCGGCCAATGAC | 9673269 |
| CYP2C23_control | AATCTCCTTTTGCATTTT | 9673293 |
